# Methylglyoxal is an antibacterial effector produced by macrophages during infection

**DOI:** 10.1101/2024.11.03.621721

**Authors:** Andrea Anaya-Sanchez, Samuel B. Berry, Scott Espich, Alex Zilinskas, Phuong M. Tran, Carolina Agudelo, Helia Samani, K. Heran Darwin, Daniel A. Portnoy, Sarah A. Stanley

## Abstract

Infected macrophages transition into aerobic glycolysis, a metabolic program crucial for control of bacterial infection. However, antimicrobial mechanisms supported by aerobic glycolysis are unclear. Methylglyoxal is a highly toxic aldehyde that modifies proteins and DNA and is produced as a side-product of glycolysis. Here we show that despite the toxicity of this aldehyde, infected macrophages generate high levels of methylglyoxal during aerobic glycolysis while downregulating the detoxification system. We use targeted mutations in mice to modulate methylglyoxal generation and show that reducing methylglyoxal production by the host promotes survival of *Listeria monocytogenes* and *Mycobacterium tuberculosis*, whereas increasing methylglyoxal levels improves control of bacterial infection. Furthermore, we show that bacteria that are unable to detoxify methylglyoxal are avirulent and experience up to 1000-fold greater genomic mutation frequency during infection. Taken together, these results suggest that methylglyoxal is an antimicrobial innate immune effector that defends the host against bacterial pathogens.

## Main text

During bacterial infection, macrophage activation relies upon a transition to aerobic glycolysis, a metabolic program in which cells produce ATP by increased flux through glycolysis rather than through oxidative phosphorylation, even in the presence of oxygen^1–3^. Increased flux through glycolysis promotes macrophage activation in part by stabilization of the transcription factor HIF-1α, which promotes transcription of inflammatory cytokine genes, most prominently IL-1β^4^. HIF-1α also promotes transcription of metabolic genes that support increased flux through glycolysis, including hexokinase-2 (HK2), an inducible isoform of hexokinase (HK)^5,6^. HIF-1α/aerobic glycolysis also mediates cell intrinsic control of intracellular bacterial pathogens^3^. Although induction of inducible nitric oxide synthase and production of nitric oxide (NO) is promoted by this metabolic program^7^, other antimicrobial effects of this pathway have not been fully elucidated. Previously we showed that IFN-γ-dependent control of *Mycobacterium tuberculosis* (*Mtb*) requires positive feedback signaling loop involving HIF-1α, aerobic glycolysis, and NO^7^. Although NO is thought to be directly antimicrobial, its ability to promote HIF-1α stabilization and aerobic glycolysis, as well as other signaling pathways during infection, leaves the nature of aerobic glycolysis-dependent antimicrobial responses uncertain.

Methylglyoxal (MG) is a highly reactive aldehyde side-product of glycolysis that damages proteins and DNA^8^. All known cells possess a MG detoxifying glyoxalase system consisting of glyoxalase I (GLO1 in mammals; GloA in bacteria) and glyoxalase II (GLO2 in mammals; GloB in bacteria)^9,10^. We previously found that deletion of *gloA* in the intracellular pathogen *Listeria monocytogenes* (*ΔgloA^Lm^*) results in increased sensitivity to MG and severe attenuation during an intravenous murine model of infection^9^, suggesting that MG detoxification plays a critical role during infection. However, the source of MG impacting *L. monocytogenes* was unknown. Here, we demonstrate that *gloA* in both *L. monocytogenes* and *Mtb* is required for defense against MG produced by host cells undergoing aerobic glycolysis. Increasing MG production by host cells results in enhanced restriction of *L. monocytogenes*, *Francisella tularensis* live vaccine strain (LVS), methicillin resistant *Staphylococcus aureus* (MRSA), and both *Mtb* and *L. monocytogenes gloA* mutants. Finally, we show that bacterial *gloA* is required for defense against mutational stress imposed by MG *in vivo*. Taken together, these data support the hypothesis that MG produced during aerobic glycolysis is a previously unappreciated innate immune antimicrobial effector. Because of its non-specific reactivity, we propose harnessing MG for antimicrobial immunity is a strategy akin to the use by immune cells of reactive oxygen and nitrogen species for controlling infection.

### Activated macrophages produce elevated levels of methylglyoxal

To evaluate whether activation of macrophages results in increased MG production^3^, we stimulated wild-type (WT) C57BL/6 bone-marrow derived macrophages (BMDMs) with the TLR2 agonist PAM3CSK4, IFN-γ, or the combination, and assessed MG levels in cell lysates (Figure 1A and Extended Data Figure 1A). Whereas either IFN-γ or PAM3CSK4 treatment increased MG production, the combination resulted in the largest increase (Figure 1A and Extended Data Figure 1A). Next, to correlate glycolytic flux with MG production, we measured glucose consumption from the media as a proxy for glycolytic flux. As expected^3^, levels of glycolysis in activated macrophages mirrored the increase in MG production observed (Figure 1B). To determine whether activated BMDMs respond to increased MG production by upregulating the glyoxalase system, we measured glyoxalase 1 (*Glo1*) mRNA. We found that although there is increased glycolytic flux in stimulated macrophages (Figure 1A), instead of upregulating *Glo-1*, activated macrophages downregulated expression of this enzyme (Figure 1C), similar to what has previously been reported in IFN-γ and LPS-activated macrophages^11^.

**Figure 1.**
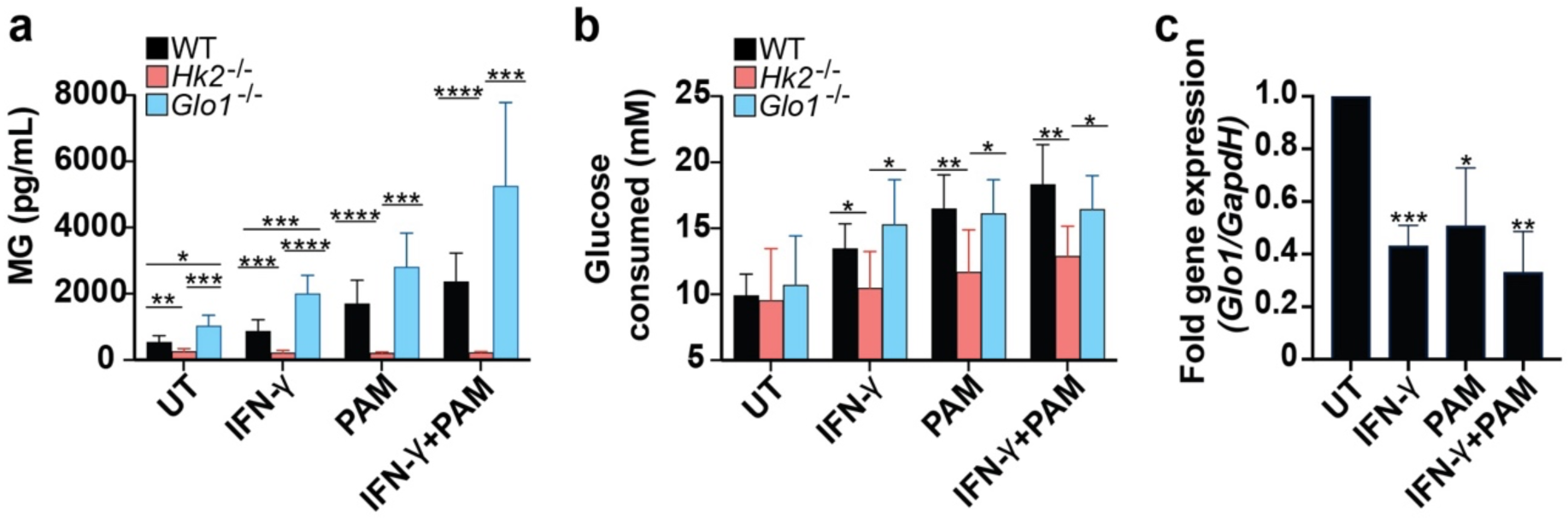
Activated macrophages produce elevated levels of methylglyoxal. **A)** Methylglyoxal measurements with an ELISA kit from Biomatik in uninfected wild-type (WT), *Hk2^fl/fl^LysMcre* (*Hk2*−/−) and *Glo1−/−* B6 murine macrophages stimulated for 24 hours with IFN-γ or PAM3CSK4 (PAM) or both. **(B)** Measurement of glucose uptake in uninfected wild-type (WT), *Hk2^fl/fl^LysMcre* (*Hk2*−/−) and *Glo1−/−* B6 murine macrophages stimulated for 24 hours with IFN-γ or PAM3CSK4 (PAM) or both. **(C)** qPCR data showing fold induction of *Glo1* in wild-type B6 macrophages that were stimulated for 24 hours with IFN-γ or PAM3CSK4 (PAM) or both. **(A)** and **(B)** were performed in parallel. Data shown are from at least three independent experiments. *P* values were calculated using Unpaired t-test; \**P* < 0.05, \*\**P* < 0.01, \*\*\**P* < 0.001, **** indicates *P* <0.0001.

We next tested whether limiting flux through glycolysis in activated macrophages results in decreased MG production. Hexokinase (HK) catalyze the conversion of glucose into glucose-6-phosphate, the first step in glycolysis. Hexokinase 2 (HK2) is an inducible isoform of HK that is not expressed at baseline in macrophages but is upregulated in activated macrophages to support enhanced glycolytic flux^5,6^. First, we tested whether deleting *Hk2* from macrophages impacted the production of MG. BMDMs from *Hk2^fl/fl^LysMcre* mice were stimulated with PAM3CSK4, IFN-γ, or both and glucose uptake was measured at 24h post stimulation. Prior to stimulation, *Hk2^fl/fl^LysMcre* deficient macrophages consumed equivalent amounts of glucose as WT (Figure 1B). However, *Hk2* deficient macrophages did not increase production of MG upon stimulation, with PAM3CSK4, IFN-γ or the combination of both (Figure 1A), consistent with a defect in upregulating aerobic glycolysis.

To increase MG production by macrophages, we generated mice lacking the MG detoxification enzyme *Glo-1* using CRISPR mediated mutation^12^. The resulting *Glo1^−/−^*mice had a 10bp deletion removing nucleotides 432-441 from exon 1 of the *Glo1* gene (Extended Data Figure 1B). To confirm that this mutation resulted in a loss of GLO-1 expression in macrophages, we prepared BMDMs from these mice, and performed qPCR in unstimulated macrophages. We found that *Glo1* transcript was undetectable in these cells (Extended Data Figure 1C). *Glo1^−/−^*BMDMs exhibited elevated MG levels compared to B6 macrophages in all conditions examined, even in unstimulated controls (Figure 1A). This excess MG production resulted in no viability defects (Extended Data Figure 1D). Importantly, *Glo1^−/−^* BMDMs had comparable increases in glucose consumption as WT B6 macrophages (Figure 1B). This demonstrates that deletion of *Glo1* does not affect flux through glycolysis, allowing us to separate the effects of increased MG production from increased glycolytic flux. Taken together, these results implicate aerobic glycolysis as the primary contributor to MG production in activated BMDMs and the glyoxalase pathway as a primary MG detoxification system.

### Modulation of host MG levels influences intracellular growth of *L. monocytogenes*

We previously showed that while *L. monocytogenes* lacking *gloA* (*ΔgloA^Lm^*), the first enzyme in the glyoxalase pathway, grows normally in broth, it has severe defects during murine infection^9^. Given that activated macrophages not only increase MG production but also down-regulate expression of their own glyoxalase enzymes (Figure 1), we hypothesized that *L. monocytogenes gloA* is important for countering host-derived MG in macrophages. To explore this hypothesis, we first tested whether MG production in BMDMs infected with *L. monocytogenes* mirrors what we previously observed with PAM3CSK4 and IFN-γ (Figure 1). Infection alone induced a modest increase in MG concentrations (Figure 2A). Activation with IFN-γ or with IFN-γ and PAM3CSK4 prior to *L. monocytogenes* infection increased MG levels (Figure 2A). As expected, *Glo1^−/−^* macrophages infected with *L. monocytogenes* exhibited elevated levels of MG in both resting macrophages, and even higher levels with activation. Conversely, *Hk2^−/−^* macrophages did not increase MG production under any conditions examined (Figure 2A). Resident peritoneal macrophages (PMs) are a primary cell type that display a more activated phenotype compared with BMDMs^13^, and have enhanced glycolytic flux^14^. Accordingly, when infected with *L. monocytogenes,* PMs had higher MG levels than BMDMs (Figure 2A).

**Figure 2.**
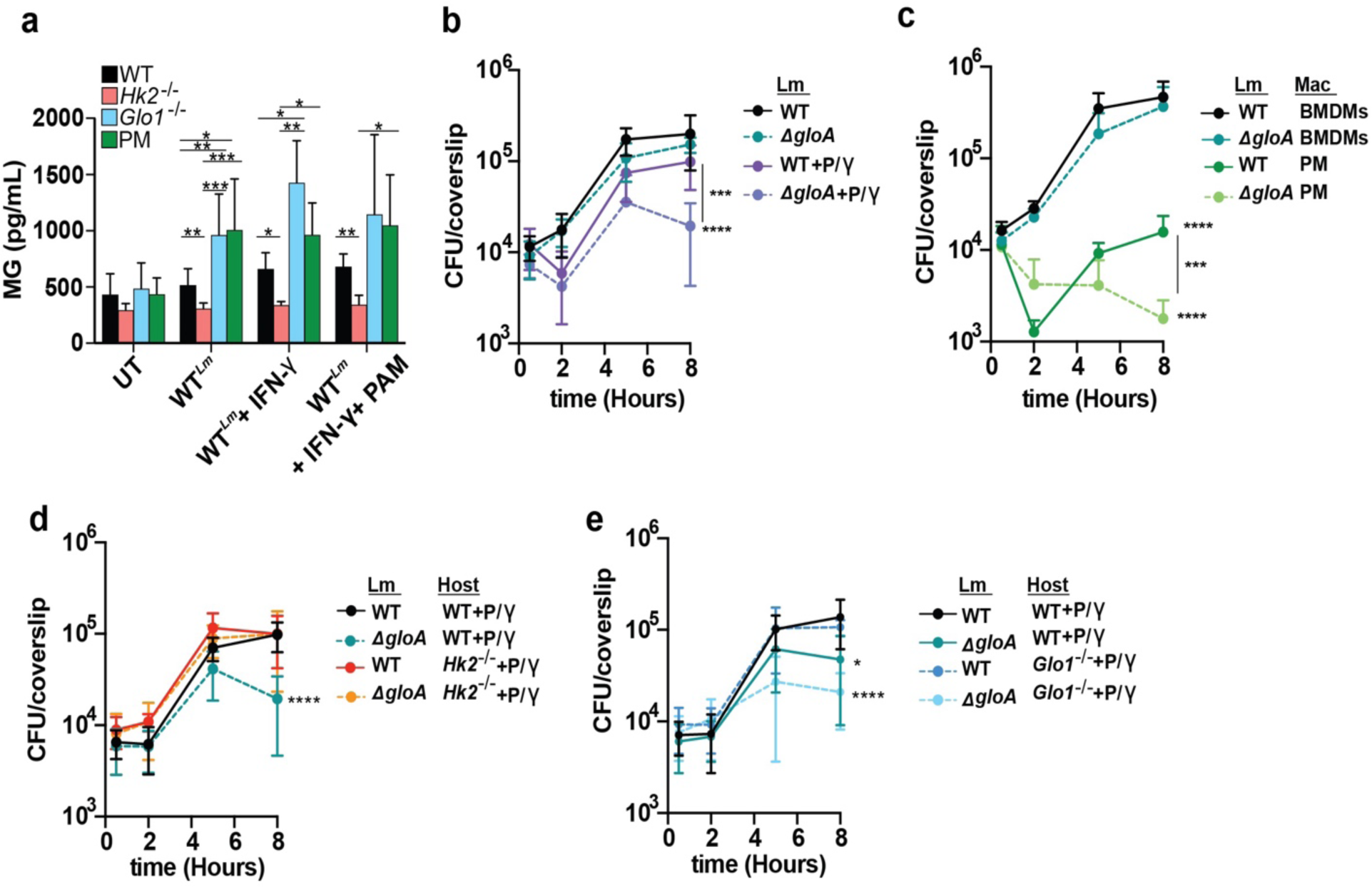
Modulation of host MG influences intracellular growth of *L. monocytogenes*. **(A)** Methylglyoxal concentrations by ELISA kit from Biomatik in wild-type (WT), *Hk2^fl/fl^LysMcre* (*Hk2*−/−) and *Glo1−/−* B6 murine macrophages that were infected with wild-type *L. monocytogenes* or stimulated for 8 hours with IFN-γ or IFN-γ and PAM3CSK4 (PAM). Data represent at least three independent experiments. WT **(B),** *Hk2^fl/fl^LysMcre* (*Hk2*−/−) **(D)** or *Glo1−/−* **(E)** BMDMs were infected with wild-type *L. monocytogenes* or the *gloA* mutant without treatment or treated with IFN-γ and PAM3CSK4 (+P/γ) and intracellular CFU were enumerated at different time-points post-infection. **(C)** WT BMDMs or peritoneal macrophages (PM) were infected with wild-type *L. monocytogenes* or the *gloA* mutant without treatment and intracellular CFU were enumerated at different time-points post-infection. **(B-E)** Data are mean and SEM of three technical replicates of three independent experiments. Statistical significance is shown for *ΔgloA* infections compared to wild-type *L. monocytogenes* growing in wild-type macrophages. *P* values were calculated using **(A)** Unpaired t-test **(B-E)** Mann-Whitney test; \**P* < 0.05, \*\**P* < 0.01, \*\*\**P* < 0.001, **** indicates *P* <0.0001.

Having established macrophage infection conditions that result in varying levels of MG, we next tested whether virulence defects of the Δ*gloA^Lm^* strain correlate with MG concentrations in macrophages by enumerating colony-forming units (CFU) of *L. monocytogenes* at multiple timepoints after infection. No difference in CFUs was observed between WT *L. monocytogenes* (WT*^Lm^*) and Δ*gloA^Lm^* at any time point post-infection of unstimulated BMDMs, consistent with the relative lack of MG produced under these conditions (Figure 2A and 2B). However, in IFN-γ or PAM3CSK4 activated macrophages, Δ*gloA^Lm^*mutants were attenuated for growth relative to WT*^Lm^* (Extended Data Figure 2A and 2B). This growth defect was exacerbated when macrophages were stimulated with both IFN-γ and PAM3CSK4 (Figure 2B). Consistent with their higher levels of MG, PMs had significantly higher bactericidal activity against WT*^Lm^* compared with BMDMs (Figure 2C). Furthermore, Δ*gloA^Lm^* was attenuated in PM even in the absence of IFN-γ (Figure 2C).

We next hypothesized that abrogating MG production during infection would rescue the phenotype of the Δ*gloA^Lm^* mutant, whereas increasing MG levels in the host would further attenuate its survival. Consistent with this hypothesis, the intracellular growth defect observed for the Δ*gloA^Lm^*mutant in IFN-γ and PAM3CSK4 activated macrophages was completely reversed to WT*^Lm^*-survival in *Hk2*^−/−^ BMDMs (Figure 2D). Conversely, in *Glo1^−/−^* knockout macrophages, Δ*gloA^Lm^* mutants had an intracellular growth defect at eight hours post-infection compared to WT bacteria even in the absence of IFN-γ, which aligns with the elevated levels of MG observed even in resting cells (Figure 2E). The Δ*gloA^Lm^* bacteria were even further attenuated for growth in IFN-γ and PAM3CSK4 activated *Glo1^−/−^*cells (Figure 2E). The growth of WT*^Lm^* was indistinguishable in WT and *Glo1^−/−^* macrophages, suggesting that bacterial GloA efficiently detoxifies even elevated levels of MG produced by macrophages. Taken together, these data suggests that MG detoxification is important for intracellular growth of *L. monocytogenes* and that modulation of MG production in the host can influence the outcome of infection of *ΔgloA^Lm^* bacteria. These results further underscore the importance of detoxification of MG for bacterial virulence.

### Host-produced MG controls infection with *L. monocytogenes in vivo*

We next sought to characterize the influence of MG production by the host during infection *in vivo*. Having demonstrated that IFN-γ promoted MG production by macrophages and suppression of the growth of a Δ*gloA^Lm^* strain, we predicted that IFN-γ-deficient mice would not restrict Δ*gloA^Lm^* growth. WT and *Ifng^−/−^* mice were intravenously infected with *L. monocytogenes* and bacterial loads in livers and spleens were measured at two days post-infection. As expected ^9^, there were approximately 1,000-fold fewer Δ*gloA^Lm^* in the livers and spleens of WT mice when compared with WT*^Lm^*-infected mice (Figure 3A and 3B). Complementation with the *gloA* gene on an integrative plasmid in the Δ*gloA^Lm^* mutant restored growth and survival (Extended Data Figure 3). *Ifng^−/−^* mice were modestly more susceptible to WT*^Lm^*, with increases in bacterial burdens of ∼10-fold in both organs. Strikingly, the virulence defect of the Δ*gloA^Lm^* mutant was almost completely reversed in *Ifng*^−/−^ mice (Figure 3A, 3B). Taken together, these data demonstrate that IFN-γ is crucial for mice to restrict growth of a Δ*gloA^Lm^*mutant.

**Figure 3.**
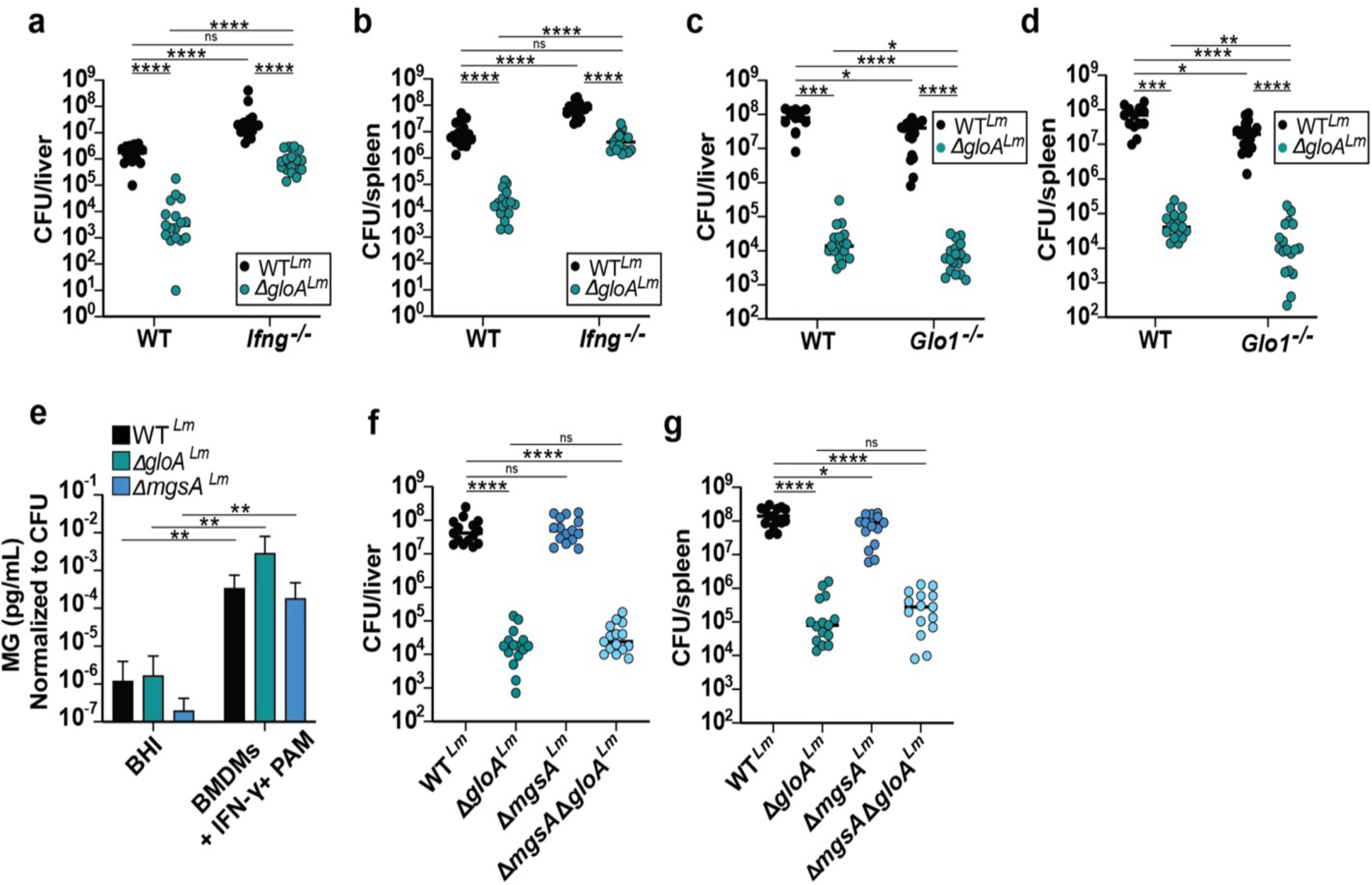
Host-produced MG controls infection with *L. monocytogenes*. **(A** and **B)** Wild-type or *Ifng−/−* B6 mice were infected with 10^4^ CFU of wild-type or *ΔgloA L. monocytogenes.* Livers **(A)** and spleens **(B)** were harvested 48-hours post-infection and CFU were enumerated. Data and median represent at least three pooled experiments. **(C** and **D)** Wild-type or *Glo1−/−* B6 mice were infected with 10^5^ CFU of wild-type or *ΔgloA L. monocytogenes.* CFU of livers **(C)** and spleens **(D)** were counted 48-hours post-infection. Data and median represent at least three pooled experiments. **(E)** Intracellular methylglyoxal concentrations from bacteria grown in BHI broth or inside of wild-type macrophages that were pre-stimulated with IFN-γ and PAM3CSK4 (PAM). Methylglyoxal concentrations were normalized to bacterial CFU counts at the time of collection. Data represent at least three independent experiments. **(F** and **G)** Wild-type B6 mice were infected with 10^5^ CFU of wild-type, *ΔgloA*, *ΔmgsA or ΔmgsAΔgloA L. monocytogenes* and CFU of livers **(C)** and spleens **(D)** were counted 48-hours post-infection. Data and median represent at least three pooled experiments. For all experiments *P* values were calculated using Mann-Whitney test; \**P* < 0.05, \*\**P* < 0.01, \*\*\**P* < 0.001, **** indicates *P* <0.0001.

We next tested whether increasing MG levels in the host enhances control of *L. monocytogenes* during infection. *Glo1^−/−^*mice had an enhanced ability to control infection of Δ*gloA^Lm^* mutants (Figure 2E), similar to what we observed in BMDMs. Interestingly, we also observed enhanced control of WT*^Lm^* in *Glo1^−/−^* mice. There was ∼10-fold fewer bacteria of both strains in livers and spleens of *Glo1^−/−^*mice when compared to WT mice (Figure 3C and 3D). Together, our findings show that detoxification of MG is essential for *L. monocytogenes* virulence *in vivo* and that increasing MG levels in the host enhances control of *L. monocytogenes* infection.

In bacteria, MG is produced as a side-product of glycolysis by the action of methylglyoxal synthase (MgsA)^10^, which converts dihydroxyacetone phosphate (DHAP) to MG. To test whether a Δ*gloA^Lm^* strain is sensitive to MgsA-derived MG, we generated an in-frame deletion of *mgsA.* In broth, deletion of *mgsA* resulted in lower MG levels in both Δ*gloA^Lm^* and WT*^Lm^* strains, consistent with disrupted MG generation from bacterial glycolysis (Figure 3E). Interestingly, bacteria growing inside of BMDMs had elevated MG levels compared to bacteria growing in broth (Figure 3E). Importantly, deleting *mgsA* had no detectable impact on intracellular MG levels during macrophage infection (Figure 3E). In mice, deletion of *mgsA* had no impact on bacterial burdens in the liver and only resulted in a slight decrease in CFU relative to WT bacteria in the spleen (Figure 3F and G). Furthermore, deletion of *mgsA* did not rescue attenuation of the Δ*gloA^Lm^* mutant (Figure 3F and 3G). Together these data support the hypothesis that a major source of MG encountered by bacteria during infection is most likely host derived.

### MG detoxification is important for *Mycobacterium tuberculosis* both *in vitro* and *in vivo*

We next hypothesized that other intracellular pathogens must also detoxify MG to survive *in vivo* and tested this hypothesis using *M. tuberculosis (Mtb),* a highly host-adapted intracellular pathogen. In broth, MG treatment of the H37Rv WT strain (WT*^Mtb^*) resulted in dose-dependent killing (Figure 4A). We next sought to identify *Mtb gloA* homologs and found four putative GloA enzymes and two putative GloB enzymes in *Mtb* (Extended Data Figure 4). It was previously suggested that Rv0911 might function as a glyoxalase in *Mtb*, however whether Rv0911 participates in MG detoxification has not been demonstrated^15^. We found that an Rv0911::tn mutant was significantly more susceptible than WT*^Mtb^* to MG, a defect that was fully restored by expression of Rv0911 using its native promoter (Figure 4A). Growth of this mutant was indistinguishable from WT*^Mtb^* in both standard 7H9 and in minimal media (Figure 4B). To assess the role of MG detoxification in *Mtb* macrophage survival, we evaluated growth of WT*^Mtb^* and Rv0911::tn in BMDMs. In contrast to what we observed with the *L. monocytogenes gloA* mutant, Rv0911::tn was attenuated for growth compared to WT*^Mtb^* even in resting macrophages. This attenuation was fully restored in the complemented strain (Figure 4C). As expected, we observed a decrease in CFU of both WT*^Mtb^* and Rv0911::tn from IFN-γ treated macrophages, with Rv0911::tn mutants being more attenuated than WT*^Mtb^* (Figure 4C). We did not observe differences in the ability of *Glo1*^−/−^ macrophages to control infection with WT*^Mtb^*, either in the presence or absence of IFN-γ. However, similar to what we observed with *L. monocytogenes*, *Glo1*^−/−^ macrophages were better able to control infection with the Rv0911::tn mutant, both in resting and IFN-γ activated macrophages (Figure 4C).

**Figure 4.**
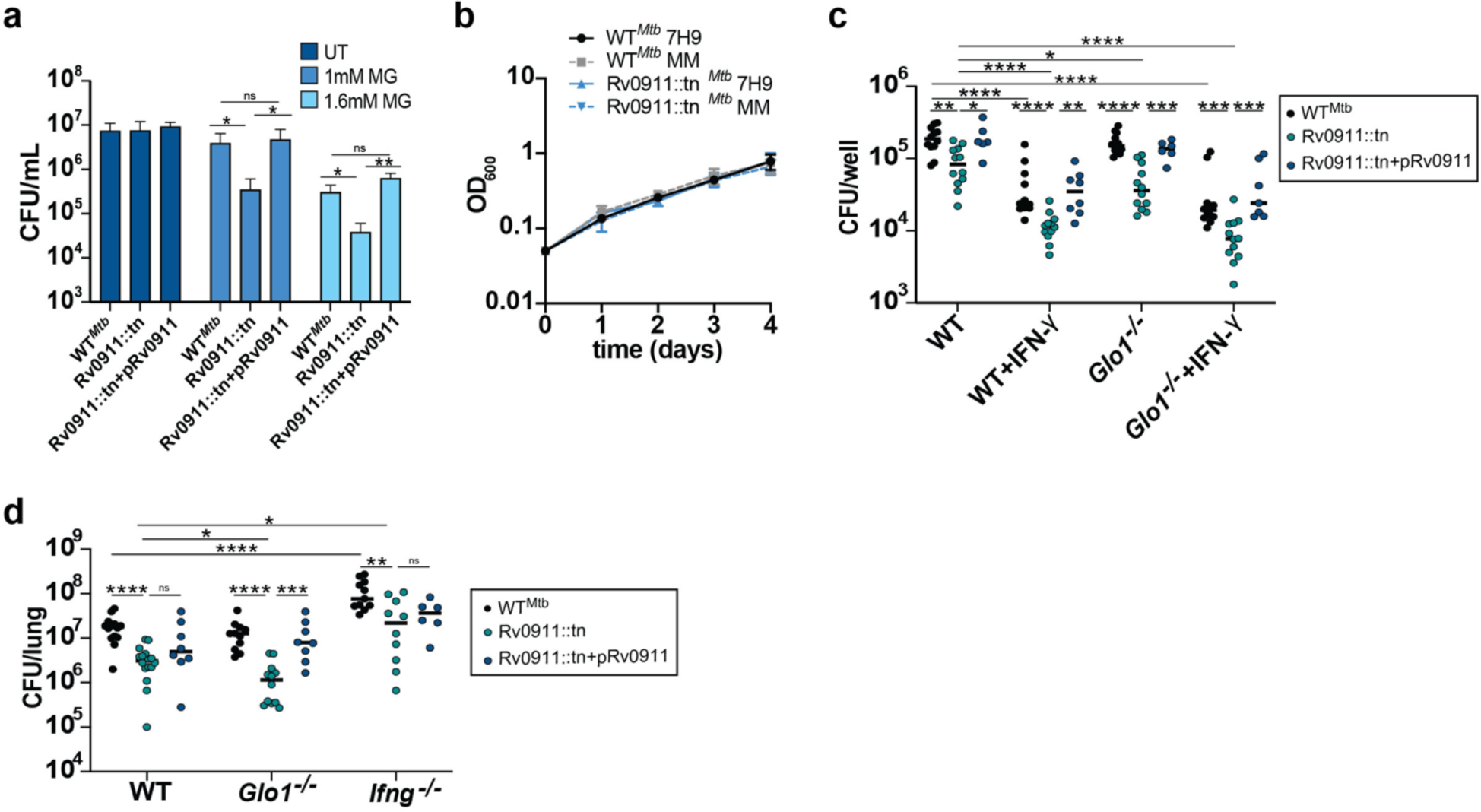
MG detoxification is important in *Mycobacterium tuberculosis*. **(A)** *Mtb* wild-type (H37Rv), Rv0911::tn or the complemented strains were treated with 1 or 1.6mM of methylglyoxal (MG) and CFU was determined after six days of exposure. Data from three independent experiments. **(B)** Growth of wild-type and Rv0911::tn mutant of *Mtb* in 7H9 and minimal media. **(C)** Wild-type and *Glo1−/−* B6 BMDMs were infected with wild-type, the Rv0911::tn mutant or the complemented strain, without treatment or treated with IFN-γ. Six days post-infection, intracellular CFU were enumerated. Data represent at least three pooled experiments. **(D)** Wild-type, *Glo1−/−* and *Ifng−/−* B6 mice were aerosol-infected with 250 CFU of wild-type (H37Rv), Rv0911::tn or the complemented strain. Lung CFU were determined at twenty-eight days post-infection. Data and median represent at least three independent experiments. For all experiments *P* values were calculated using **(A)** Unpaired t-test, **(B-D)** Mann-Whitney test; \**P* < 0.05, \*\**P* < 0.01, \*\*\**P* < 0.001, **** indicates *P* <0.0001.

To determine the importance of GloA for *Mtb* infection *in vivo*, we performed aerosol infections of mice. At twenty-eight days post-infection of WT mice, we observed ∼10-fold reduction of Rv0911::tn CFU compared to WT*^Mtb^*, a defect that was restored with complementation (Figure 4D). As observed with *L. monocytogenes*, the survival of Rv0911::tn mutants was partially restored in *Ifng*^−/−^ mice (Figure 4D). In *Glo1^−/−^* deficient mice we did not observe enhanced control of infection of WT*^Mtb^*, but the Rv0911::tn mutant was significantly more attenuated than in WT mice (Figure 4D). Thus, the attenuation of the Rv0911::tn mutant was partially reversed in the absence of IFN-γ dependent immunity *in vivo*, and enhancing MG levels by the host improves control of a *Mtb* Rv0911::tn mutant. These results mirror those obtained from our experiments with *L. monocytogenes*, suggesting that MG production by the host is an innate immune response that must be countered by multiple bacterial pathogens during infection.

### MG sensitivity varies widely across bacterial species

Both Mtb and *L. monocytogenes* are relatively resistant to MG in the presence of an intact glyoxalase system. To evaluate MG sensitivity among other bacterial genera, we exposed a panel of gram-positive and gram-negative bacteria to a range of MG concentrations and assessed survival. We observed a large range of susceptibility to MG across the species tested, suggesting that bacteria possess different innate capacities to detoxify MG (Figure 5A). Interestingly, we found that the live vaccine strain of *Francisella tularensis* (FtLVS) exhibited enhanced susceptibility to MG than other bacterial pathogens tested. To determine whether the relatively high susceptibility of FtLVS to MG *in vitro* translates to enhanced sensitivity to host derived MG, we infected WT and *Glo1^−/−^* mice with FtLVS and measured CFU in lungs five days after infection. The bacterial burden of FtLVS was dramatically reduced in *Glo1^−/−^*mice compared to WT mice (Figure 5B). Methicillin resistant *Staphylococcus aureus* (MRSA) is an extracellular pathogen; however, restriction of this pathogen also requires the action of macrophages^16^. MRSA exhibited an intermediate susceptibility to MG in axenic culture (Figure 5A). We infected WT and *Glo1^−/−^* mice with MRSA and enumerated CFU three days post-infection. We observed a six-fold attenuation in *Glo1^−/−^* mice compared to WT mice (Figure 5C). Taken together, these data demonstrated that host derived MG is an innate immune effector that can help control infection with multiple bacterial pathogens. How much MG they encounter might depend on the inflammatory response to each pathogen; whereas how toxic MG is to the bacteria might depend on the efficiency of MG detoxification by bacteria.

**Figure 5.**
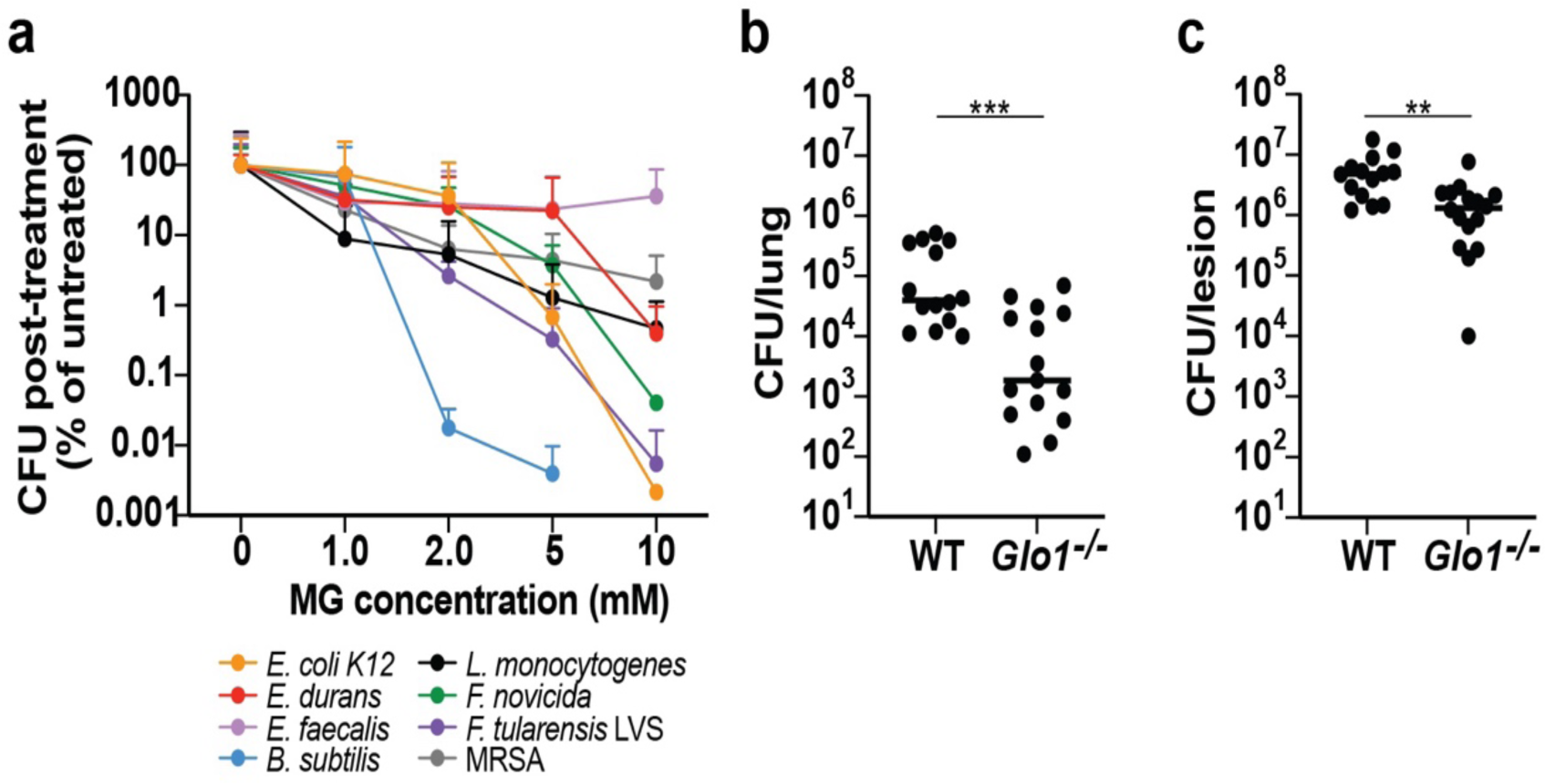
MG detoxification is important for many pathogens. **(A)** Survival panel of *Escherichi coli K12, Enterococcus durans, Enterococcus faecalis, Bacillus subtilis, Listeria monocytogenes, Francisella novicida, Francisella tularensis live-vaccine strain (*LVS) and methicillin-resistant *Staphyloccocus aureus (*MRSA) grown to mid-log (0.5-1 OD_600_) and exposed to various methylglyoxal concentrations for 1 hour. Data are normalized to the untreated condition and represent at least two independent experiments. **(B)** Lung CFU of wild-type and *Glo1−/−* mice infected with 10^4^ CFU of *F. tularensis LVS*. Data and median represent at least three pooled experiments. **(C)** CFU of skin lesions dissected from WT B6 and *Glo1*−/− mice that were subcutaneously infected with 1×107 CFU WT MRSA strain JE2 USA300. For all experiments *P* values were calculated using Mann-Whitney test; \*\**P* < 0.05, \*\**P* < 0.01, \*\*\**P* < 0.001, **** indicates *P* <0.0001.

### MG detoxification prevents bacterial mutations during infection

MG is a highly reactive electrophile that reacts with guanine bases in DNA, resulting in damage and potential mutagenesis^17^. We previously reported that exposure of *L. monocytogenes* to MG in broth increased mutation frequency, a phenotype that is exacerbated in bacteria lacking GloA^9^. Accordingly, we next assessed whether bacteria experience an enhanced mutational burden during infection using a rifampicin resistance assay. Resistance to the antibiotic rifampicin results from the acquisition of point mutations within its target, bacterial RNA polymerase ^18^. Thus, the number of rifampicin mutants in a population of bacteria is a proxy for estimating mutation rates. To determine the mutation rate of *L. monocytogenes in vivo*, we performed IV mouse infections and inoculated bacteria harvested from livers and spleens on plates with and without rifampicin. The Δ*gloA^Lm^* mutant had over 100-fold higher mutation frequency compared to WT*^Lm^* (Figure 6A and B). The increased mutation frequency was restored in a Δ*gloA^Lm^* complemented strain (Extended Data Figure 5A). Thus, bacterial detoxification of MG produced *in vivo* protects the genome from increased mutation rates. To test the hypothesis that host derived MG production causes bacterial DNA damage, we determined mutation frequency in *Glo1^−/−^* and *Ifng^−/−^* mice. As predicted, the mutation frequency for both WT*^Lm^* and Δ*gloA^Lm^* strains increased in the liver and spleen during infection of *Glo1^−/−^* mice that have higher concentrations of MG compared to WT mice (Figure 6A and B). Conversely, the difference in mutation frequency between WT*^Lm^* and Δ*gloA^Lm^* was reduced in *Ifng^−/−^* mice (Figure 6C and D). To further understand the observed increase in mutation frequency in mutants lacking GloA*^Lm^*, we followed *L. monocytogenes* CFU and mutations frequency over time during an IV mouse infection. We observed that the growth kinetics of Δ*gloA^Lm^*were similar to the WT*^Lm^* for the first 12 hours, at which time the attenuated growth and increased mutation frequency phenotypes of the mutant emerged simultaneously (Figure 6E and F; Extended Data Figure 5C and 5D). Finally, given that GloA*-*dependent virulence phenotypes in *L. monocytogenes* were recapitulated in *Mtb,* we tested whether *Mtb* also accumulates increased DNA damage when its glyoxalase pathway is disrupted. To test this hypothesis, we infected mice with the Rv0911::tn strain and determined the frequency of emergence of rifampicin resistant colonies. We observed that the Rv0911::tn mutant had a 10-fold higher mutation frequency compared to WT*^Mtb^*(Figure 6G). The mutation frequency was restored to WT levels in a Rv0911::tn complemented strain (Extended Data Figure 5B). Mutation frequency experienced by the Rv0911::tn mutant was higher in *Glo1^−/−^* mice and restored in *Ifng^−/−^*mice (Figure 6G). Although the mutation frequency of WT*^Mtb^* was not impacted by *Glo1* deficiency, the mutation rate was decreased in the absence of IFN-γ (Figure 6G). Together these data demonstrate that MG detoxification is important for DNA protection of both *L. monocytogenes* and *Mtb,* and that IFN-γ dependent immunity is required for MG-dependent mutagenesis of bacterial genomes.

**Figure 6.**
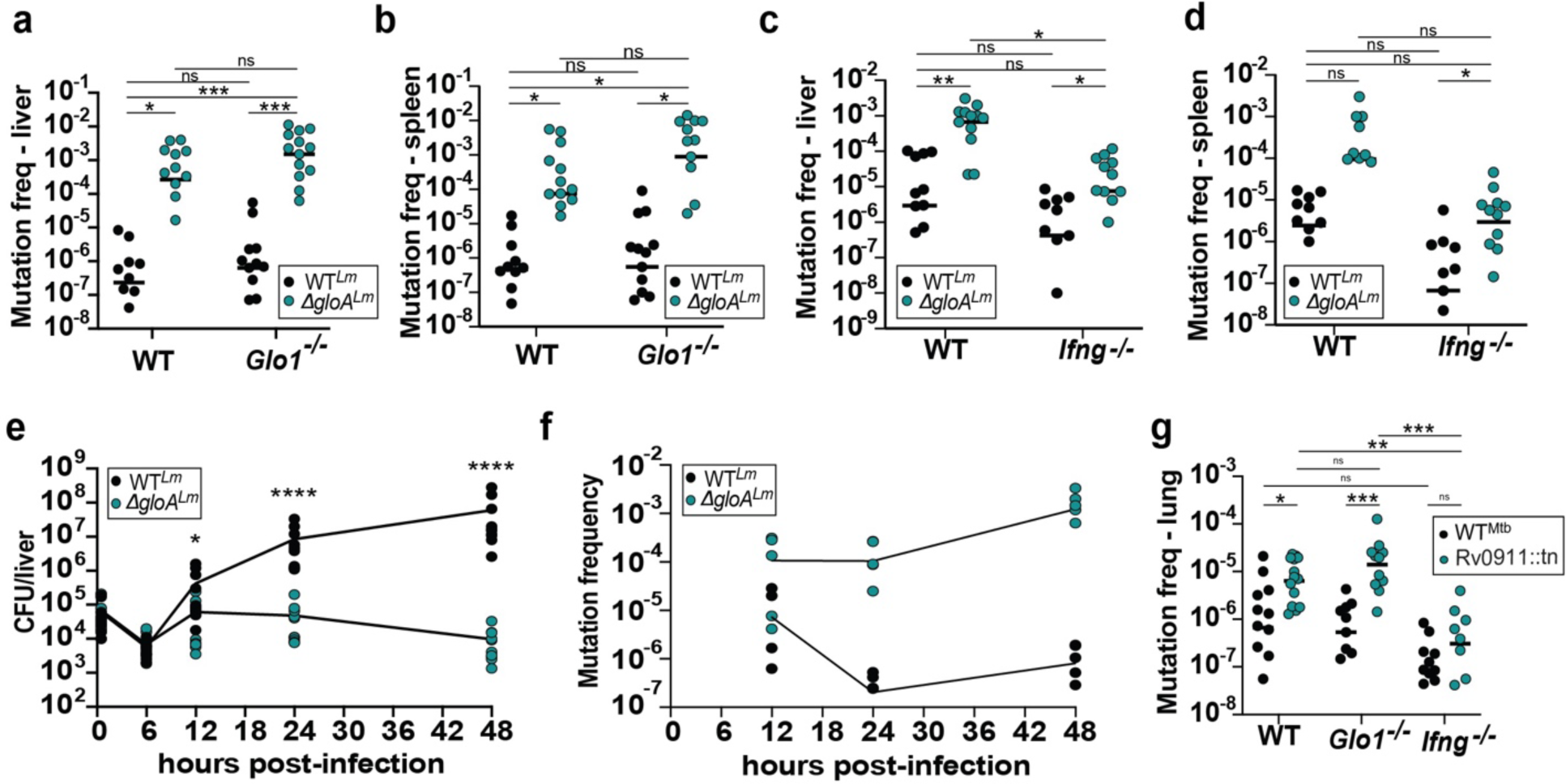
MG detoxification is important for bacterial DNA protection. **(A** and **B)** Wild-type (WT) and *Glo1−/−* B6 mice were infected with 10^5^ CFU of wild-type or *ΔgloA L. monocytogenes.* Livers **(A)** and spleens **(B)** were harvested 48-hours post-infection and whole organs were plated on agar plates containing rifampicin. **(C** and **D)** Wild-type (WT) and *Ifng−/−* B6 mice were infected with 10^4^ CFU of wild-type or *ΔgloA L. monocytogenes.* Livers **(C)** and spleens **(D)** were harvested 48-hours post-infection and whole organs were plated in agar plates containing rifampicin. **(E)** Wild-type (WT), *Glo1−/−* and *Ifng−/−* B6 mice were aerosol-infected with 250 CFU of wild-type (H37Rv), Rv0911::tn or the complemented strain. Lungs were harvested at twenty-eight days post-infection and whole organs were plated on agar plates containing Rifampicin. **(F** and **G)** Wild-type B6 mice were infected with 10^5^ CFU of wild-type or *ΔgloA L. monocytogenes.* Livers were harvested at various time points post-infection and CFU **(F)** or mutation frequency **(G)** were determined for whole organs. For all experiments, mutation frequency was calculated as the total rifampicin resistant colonies normalized to total CFU per organ. Data and median represent at least three independent experiments. *P* values were calculated using Mann-Whitney test; \**P* < 0.05, \*\**P* < 0.01, \*\*\**P* < 0.001, **** indicates *P* <0.0001.

## Discussion

The goal of this study was to test the hypothesis that host-derived MG is antimicrobial during infection. We showed that activated macrophages increase MG production through induction of aerobic glycolysis, a process that is enhanced by IFN-γ, and downregulate the glyoxalase system that primarily detoxifies this toxic aldehyde^8^. Next, we demonstrated that bacterial GloA mutants in both *L. monocytogenes* and *Mtb* exhibit enhanced susceptibility to MG and are attenuated for virulence during murine infections. To link these phenotypes to host-derived MG, we showed that the virulence defects of both *gloA* mutants were exacerbated in hosts that produce excess MG (*Glo1^−/−^*) and alleviated in mice that lack IFN-γ. The bacterial enzyme MgsA, which produces MG during glycolysis in bacteria, contributes to MG levels in bacteria grown in broth but is dispensable for elevated MG levels seen in intracellular bacteria, and it does not contribute to the attenuation of GloA mutants, supporting the hypothesis that host-derived MG is the major contributor to the phenotype. Finally, we found that MG detoxification is important to prevent the accumulation of genomic mutations in both *L. monocytogenes* and *Mtb.* Together these data suggest that MG production is be part of the innate immune response to pathogens. Moreover, we showed that different pathogens are susceptible to MG and that this susceptibility is reflected *in vivo,* suggesting that pathogens encounter this stress during infection and have evolved mechanisms to counter this host defense. How MG is killing pathogens is not fully understood, but we demonstrated that detoxification of this toxic metabolite is important to prevent mutations in both *L. monocytogenes* and *Mtb*.

MG is produced as a side-product of metabolism, glycolysis being the primary generating pathway. As such, both bacteria and host cells produce this toxic metabolite as part of their central metabolism^8,10^. Given that MG is toxic, organisms need to detoxify it, and this is accomplished mainly through the action of the glyoxalase system that is ubiquitous across organisms^10^. Even though all known pathogens possess a glyoxalase system, our data showed that each bacterial species has a different capacity to withstand MG stress. Further studies to understand the expression and regulation of glyoxalase enzymes might shed some light into how different pathogens use this pathway during infection. In plants and some bacteria, MG can be detoxified by other enzymes^17,19^. It is possible that the difference in responding to MG by the bacteria assessed here could be a tied to the presence or absence of these alternative detoxification pathways. It is also possible that other factors, such as the permeability of bacterial cell wall/envelope to aldehydes, may also be important factors.

The successful establishment of *Mtb* infection largely depends on its ability to resist redox stress during its early interactions with the host innate immune cells^20^. We speculate that the very well adapted human pathogen *Mtb,* which was the only pathogen in this study that showed no difference in susceptibility between B6 and *Glo1^−/−^* mice, possesses very effective and redundant mechanisms that either detoxify MG or repair MG-induced damage. Additionally, *Mtb* induces a strong immune response to which it adapts to in its host to succeed in the host-pathogen coevolutionary arms race^21^. Unlike *Mtb,* we showed that the live-vaccine strain of *F. tularensis* (FtLVS) was the most attenuated WT bacteria in *Glo1^−/−^* mice compared to B6 mice. *F. tularensis* has various mechanisms to evade immune recognition and infection by this pathogen is characterized by exceptionally low inflammation at early stages of infection^20,22,23^. We hypothesize that LVS is particularly sensitive to MG and MG-induced *in vivo* stress because LVS has not evolved to resist high inflammatory settings like *Mtb*. Our lab continues to use LVS as a model to study aldehyde response and its role in pathogenesis.

It is unclear why the *ΔgloA^Lm^* mutant was more highly attenuated in vivo than the *Mtb* Rv0911::tn mutant. Unlike *Mtb*, which is an obligate human pathogen, *L. monocytogenes* is capable of transitioning from saprophyte to intracellular pathogen of mammals. Upon entry into the host cell, *L. monocytogenes* activates its virulence gene expression program by activating its master virulence regulator PrfA^24^. Activation of PrfA requires the allosteric binding of glutathione (GSH) that is produced by bacterial *gshF,* an enzyme that is robustly induced upon infection^25^. The results of our previous study showed that exposure to exogenous MG triggers *gshF* expression, suggesting that it is one of the biological and redox cues that *L. monocytogenes* senses inside the host cells to activate its virulence program. Furthermore, the GSH requirement for PrfA activation can be bypassed by mutations in PrfA (PrfA*) that lock the protein in its active conformation^25^. PrfA* mutations restore virulence of *ΔgloA^Lm^* mutants^9^. The glyoxalase system mediates MG detoxification by conjugation with GSH, which led us to hypothesize that the need to detoxify MG to activate PrfA came from the need to have available GSH^9^. Together our data suggest that the need to detoxify MG leads to upregulation of *gshF* and production of glutathione that has the dual roles of mediating MG detoxification and PrfA activation^9^. Altogether these data describe how MG is intricately involved in *L. monocytogenes* pathogenesis. In other bacteria, such as *Mtb* loss of *gloA* also reduces virulence, but not to the extent as in *L. monocytogenes*.

Although the sensitivity of GloA mutants to MG coupled with their virulence defects implicates MG in direct host antimicrobial responses, it is also possible that MG contributes to host inflammatory responses. MG reacts with arginine, cysteine, and lysine residues^26^ which results in the formation of advanced glycation end-products (AGEs) that have been associated with multiple diseases^27^. AGE recognition by receptors for advanced glycation end products (RAGEs) leads to activation of NF-kB and production of inflammatory cytokines that influence macrophage polarization^28–30^. A previous study showed that RAGE-deficient mice were more susceptible to *Mtb* infection^31^. More studies are needed to understand if MG produced by the burst of aerobic glycolysis in activated macrophages contributes to RAGE and inflammatory activation are needed.

*In vivo* mutation frequencies for *L. monocytogenes* were elevated relative to bacteria growing in broth^9^, suggesting that intracellular bacteria are exposed to mutagenic host stressors^32,33^. MG is a toxic electrophile that can react with DNA^34,35^. Our data demonstrated that MG detoxification by a glyoxalase system is necessary to prevent accumulation of mutations *in vivo*, however it is possible that MG also acts synergistically with other stressors^36–38^. While excessive acquisition of mutations can be detrimental, mutations can also be an important source of adaptation^39^. For example, *Mtb* clinical isolates often exhibit drug resistance resulting almost exclusively from the accumulation of point mutations during infection^40^. In this study we demonstrated that GloA is necessary to suppress the emergence of mutations that confer resistance to the front-line TB drug rifampicin^41^ that would otherwise be induced by IFN-γ dependent MG production. In recent years, the emergence of multidrug resistant strains of *Mtb* represent a serious health problem worldwide. Thus, how MG shapes both bacterial evolution and acquisition of drug resistance in *Mtb* and other bacterial pathogens is an important focus for future studies.

## Methods

### Ethics statement

All animal work was done in strict accordance with the recommendations in the Guide for the Care and Use of Laboratory Animals of the National Institutes of Health and the University of California, Berkeley regulations. Protocols were reviewed and approved by the Animal Care and Use Committee at the University of California, Berkeley AUP 2016-05-8811 and protocol no. R353-1113B.

### Bacterial cultures and strains

All strains used in this study are listed in Extended Data Table 1. The parental strain for all *L. monocytogenes* strains used in this study is 10403S. Bacteria were cultivated overnight at 30°C slanted in Brain-Heart Infusion (BHI; BD) with streptomycin at 200 μg/mL (GoldBio). Deletion of *mgsA* was performed by allelic exchange with the PLIM plasmid. Briefly, plasmids were constructed and transformed into XL1 Blue and SM10 *E. coli*, recovered on LB agar plates containing 100 ug/mL of carbenicillin (Sigma-Aldrich) and conjugated into *L. monocytogenes* strains. *L*. *monocytogenes* carrying the knock-out plasmid were selected on chloramphenicol at 30°C and re-streaked at 42°C three consecutive times to select for chromosomal integration. This selected strain was serially plated on DL-4-chlorophenylalanine (Acros Organics) to select for loss of PLIM. Mutants that lost the plasmid were identified by Sanger sequencing.

The *M. tuberculosis* strain H37Rv was used for all experiments. The transposon mutant Rv0911::tn was picked from an arrayed transposon mutant library generated at the Broad Institute. For genetic complementation, the pMV306 vector was used containing the open reading frame of the Rv0911 gene under the native promoter. *M. tuberculosis* strains were grown in Middlebrook 7H9 broth media supplemented with 10% OADC and 0.05% Tween 80 at 37°C. For minimal media *M. tuberculosis* growth curves, Sauton media was prepared and used. For methylglyoxal sensitivity assays, strains of *M. tuberculosis* were grown in 7H9 media buffered at pH 5.5 until mid-log (OD_600_ 0.5-0.8). Bacteria were back-diluted to OD_600_ 0.08 into acidified 7H9 media with or without methylglyoxal (MG) and grown for six days. CFU was determined by plating bacteria in 7H10 solid media.

### Mouse strains

C57BL/6J (B6, Cat #000664) mice were obtained from The Jackson Laboratory (Bar Harbor, ME) and bred in-house. *Hk2^fl/fl^* mice were obtained from the European Mutant Mouse Archive. *LysMCre* (Cat #:004781) mice were obtained from The Jackson Laboratory. *Glo1−/−* mice were generated as previously described^12^ and bred in-house. Briefly, knockout mice were generated by pronuclear injection of Cas9 mRNA, sgRNA, and a 154 nucleotide ultramer oligo (IDT Inc., Coralville, IA) containing nucleotide homology arms and encoding the *Glo1* deletion as well as synonymous mutations that disrupt sgRNA binding. Founder mice were genotyped for *Glo1* mutations by PCR using the reverse primer 5’ CCACACCTGAATGAGTCTTGCC 3’ and the forward primers 5’ GGCGTCCAGTGGCCTC 3’, 5’ CCACAGCCGGCGTCAC 3’. Mice were bred to generate homozygous lines that are maintained in-house.

### Intracellular growth curves of *L. monocytogenes*

Macrophage growth curves were performed as previously described^9,42^. Briefly, 3×10^6^ BMDMs were plated in 60 mm petri dishes containing 12 mm glass coverslips in each dish and were allowed to adhere overnight. For indicated experiments, BMDMs were stimulated for 16-18 hours with medium containing PAM3CSK4 (Invivogen) at a final concentration of 100ng/mL or recombinant mouse IFN-*γ* (R&D Systems, 485-MI) at 6.25 ng/mL. BMDMs were infected the next day at a multiplicity of infection (MOI) of 0.25 for 30 minutes, washed twice with sterile PBS and 50 μg/mL gentamicin (Sigma-Aldrich) was added 1-hour post-infection. Three coverslips were removed at each time point and plated on LB agar with streptomycin to determine CFU. Each experiment represents the average of three coverslips per time point from three independent experiments.

### *M. tuberculosis* infection of murine macrophages

BMDMs were seeded at 3×10^4^ cells per well in a 96-well TC-treated dish and were allowed to adhere overnight. For indicated experiments, BMDMs were stimulated for 16-18 hours with medium containing PAM3CSK4 (Invivogen) at a final concentration of 100ng/mL or recombinant mouse IFN-γ (R&D Systems, 485-MI) at 6.25 ng/mL. BMDMs were infected with DMEM supplemented with 5% FBS and 5% horse serum at a MOI of 1. Following Spinfection at 1200rpm for 10 minutes, infection medium was removed, and cells were washed with PBS before fresh medium was added. Fresh medium was added every two days during the infection. For CFU enumeration, six days post-infection cells were lysed in water with 0.5% Triton-X and incubated at 37°C for 10 min. Following the incubation, lysed cells were resuspended and serially diluted in PBS with 0.05% Tween-80. Dilutions were plated on 7H10 plates. CFU was determined twenty-one days after plating.

### Host cell viability assays

BMDMs were seeded at 3×10^4^ cells per well in a 96-well TC-treated dish and allowed to adhere overnight. Macrophages were either stimulated for 16-18 hours with medium containing PAM3CSK4 (Invivogen) and/or recombinant mouse IFN-*γ* (R&D Systems, 485-MI), or infected with *L. monocytogenes* strains at a MOI of 0.25 for 8 hours. To determine viability, CellTiter-Glo® Luminescent Cell Viability Assay (Cat# G7570; Promega) was used as per the manufacturer’s protocol.

### Methylglyoxal ELISA assay kit

For MG measurements in Figure 1 and 2, a MG ELISA kit (Cat# EKN53482-96T; Biomatik) that detects MG through competitive inhibition was used as per the manufacturer’s protocol. Briefly, BMDMs were seeded at 3×10^4^ cells per well in a 96-well TC-treated dish. For host MG ELISAs, supernatants combined with lysates from BMDM in 96-well plates were used. BMDMs were stimulated for 16-18 hours with PAM3CSK4 (Invivogen) at a final concentration of 100ng/mL or recombinant mouse IFN-*γ* (R&D Systems, 485-MI) and used at 6.25 ng/mL. For infected macrophage measurements, supernatants combined with lysates of BMDMs that were infected with WT *L. monocytogenes* at a MOI of 0.25 for 8 hours were used. Macrophages were lysed with 0.5% Triton-X at 37°C for 10 minutes. For bacterial measurement of MG, overnights of *L. monocytogenes* were used to infect BMDMs at MOI 1 for 30 minutes. Cells were washed with PBS and extracellular bacteria was killed with gentamycin after 1 hour. After 8 hours of infection, macrophages were lysed with 0.5% Triton-X at 37°C for 10 minutes and bacteria were transferred to a 15mL Falcon tube, washed two times in PBS and plated on LB agar with streptomycin for determining CFU. Remaining bacteria were lysed using 0.1 mm-diameter silica-zirconium beads and the lysate was used for the ELISA probe. For measurements of bacteria growing in broth, overnights of *L. monocytogenes* were back diluted in fresh BHI and grown for 8 hours. Bacteria were processed as described above. Each experiment represents the average of two technical replicates from three independent experiments. For the secondary method for MG measurements shown in Supplementary figures, a MG Fluorogenic probe (Catalog # K461-100; BioVision) that measures MG using an enzyme-coupled reaction that reduces the fluorogenic probe was used as per the manufacturer’s protocol. Samples were prepared as described above for the ELISA kit. This kit is not manufactured any more.

### Glucose assay

Glucose consumption from the media was measured using a glucose (HK) assay kit (GAHK20; Sigma-Aldrich). The protocol was modified to perform the assays in 96-well plates with 100-μl reactions as described previously^7^. Glucose consumption was calculated by measuring glucose levels in the media after infection and subtracting from glucose measured in cell-free media. Glucose and MG assays were performed in parallel.

### Mouse infections

Eight- to twelve-week-old C57BL/6J mice (Jackson Laboratory) were infected intravenously via the tail vein with 1×10^5^ CFU of *L. monocytogenes* strains in 200μL of sterile PBS as described^43^. *Ifng−/−* mice were infected with 1×10^4^ CFU. 48-hours post-infection, mice were euthanized, and spleens and livers were harvested, homogenized in 0.1% NP-40 (Sigma-Aldrich) in water, and plated on LB agar with streptomycin. Data shown represents at least three independent experiments. For infection with *M. tuberculosis,* eight- to twelve-week-old C57BL/6 mice (Jackson Laboratory) were infected with 250 CFU of *M. tuberculosis* strains by the aerosol route using a Glas-Col (Brazil, IN) full-body inhalation exposure system. Infections were allowed to proceed for twenty-eight days at which time mice were euthanized and CFU from the lungs enumerated by plating on 7H10 plates for twenty-one days. Data shown represents at least three independent experiments. For infections with the live-vaccine strain of *Francisella tularensis* (LVS), frozen stock vials were thawed and diluted into sterile PBS. Eight- to twelve-week-old C57BL/6J mice were infected subcutaneously with 100uL of inoculum with a total of 1×10^4^ CFU of LVS. Infection was allowed to proceed for five days at which time mice were euthanized and CFU from the lungs enumerated by plating on Mueller Hinton plates supplemented with FBS, 10% glucose, 2.5% ferric pyrophosphate and IsoVitalex. Colonies were counted 2 days after plating. Data shown represents at least three independent experiments. WT MRSA strain JE2 USA300 was cultivated in LB broth at 37°C for in vivo experiments. WT and *Glo1−/−* mice were subcutaneously infected on right dorsal flank between the hip and hind leg with 1×107 CFU WT MRSA strain JE2 USA300. Mice were sacrificed three days post-infection and skin lesions were dissected using 12 mm biopsy punches to standardize tissue extraction.

### MG survival assays bacterial panel

Bacterial panel experiments were performed as previously described for another aldehyde^44^. Briefly, bacteria (*Listeria monocytogenes, Enterococcus faecalis, Escherichia coli,* Methicillin-Resistant *Staphylococcus aureus, Bacillus subtilis, Enterococcus durans, Francisella tularensis LVS and Francisella novicida*) were inoculated overnight in TSBC (TSB + cysteine 0.1%) and grown at 37°C, except for LVS bacteria that were grown in Mueller Hinton broth supplemented with FBS, 10% glucose, 2.5% ferric pyrophosphate and IsoVitalex. Overnight bacterial cultures were back-diluted 1:1000 in fresh TSBC and grown to mid-log (OD_600_ of 0.5-1.0). At this point, bacteria were normalized to OD_600_ of 1, washed and resuspended in sterile PBS. LVS bacteria were not back diluted since they reach mid-log after 30 hours. Bacteria were diluted 1:100 into sterile PBS containing various concentrations of MG and incubated at 37°C for 1 hour. CFU was determined by plating on TSA supplemented with 0.1% cysteine at 37°C overnight. Data shown represents at least three independent experiments performed in parallel for all bacteria.

### Rifampicin mutagenesis assay

Mutation frequency in rifampicin was determined using similar previously described methods^18^. For rifampicin mutation frequency determination from *L. monocytogenes* after infection, 48-hours post-infection mice were euthanized, and spleens and livers were harvested, homogenized in 0.1% NP-40 (Sigma-Aldrich) in water, and plated on LB agar with streptomycin and in LB agar plates containing 5 μg/mL of rifampicin. CFUs were counted after a 72-hour incubation at 37°C. Similarly, for *M. tuberculosis* rifampicin mutation frequency measurements, lungs were harvested and plated in 7H10 plates containing 1ug/mL of rifampicin. Mutation frequency was calculated as the ratio between CFUs enumerated in the agar plates containing rifampicin and the total number of bacteria plated. Data represents the average of at least three independent experiments.

### Quantitative RT-PCR of mammalian transcripts

For quantitative RT-PCR, 3×10^5^ BMDMs were seeded in 24-well dishes and stimulated with PAM3CSK4 and IFN-*γ* as described above. At 16-18 h post-stimulation, cells were washed with PBS and lysed in 1 ml TRIzol (Invitrogen/Life Technologies). Total RNA was extracted using chloroform, and the aqueous layer was purified using RNeasy spin columns (Qiagen). cDNA was generated from 1 μg of RNA using SuperScript III (Invitrogen/Life Technologies) and oligo(dT) primers. Selected genes were analyzed using Maxima SYBR Green qPCR master mix (Thermo Scientific). Each sample was analyzed in triplicate on a CFX96 real-time PCR detection system (Bio-Rad Laboratories). Quantification cycle values were normalized to values obtained for GAPDH, and relative changes in gene expression were calculated using the ΔΔ quantification method as described before^7^.

### Statistical analysis

Data were analyzed using GraphPad Prism 10. * Indicates *P* <0.05; ** indicates *P* <0.01, *** indicates *P* <0.001, **** indicates *P* <0.0001. Statistical outliers were not excluded in this study.

## Acknowledgments

We thank Angus Y. Lee and Harmandeep S. Dhaliwal (UC Berkeley Cancer Research Laboratory) for generation of *Glo1* mutant mice. We thank Karen Elkins (FDA) for the gift of the Francisella LVS strain as well as helpful advice and protocols. Funding: We thank the NIH for the following support: 1R01AI113270-01A1 to S.A.S., 1R01AI153197-01 to S.A.S., 1P01 AI063302 to D.A.P., 1R01 AI027655 to D.A.P. and CEND Fellowship to A.A.S. We want to thank the members of the Portnoy, Stanley, Cox and Vance labs for helpful discussions. We thank Angela Hung (UCLA) for help with the mice experiments.

## Author contributions

Conceptualization and methodology, A.A.S, S.B, P.M.T, H.D, D.A.P, S.A.S; Investigation, A.A.S, S.B, S.E, A.Z, C.A, H.S; Writing-original draft, A.A.S, D.A.P, S.A.S; Writing-review & editing, S.B, P.M.T, H.D; Supervision and Funding Acquisition, D.A.P, S.A.S.

## Extended data figures

**Extended Data Figure 1.**
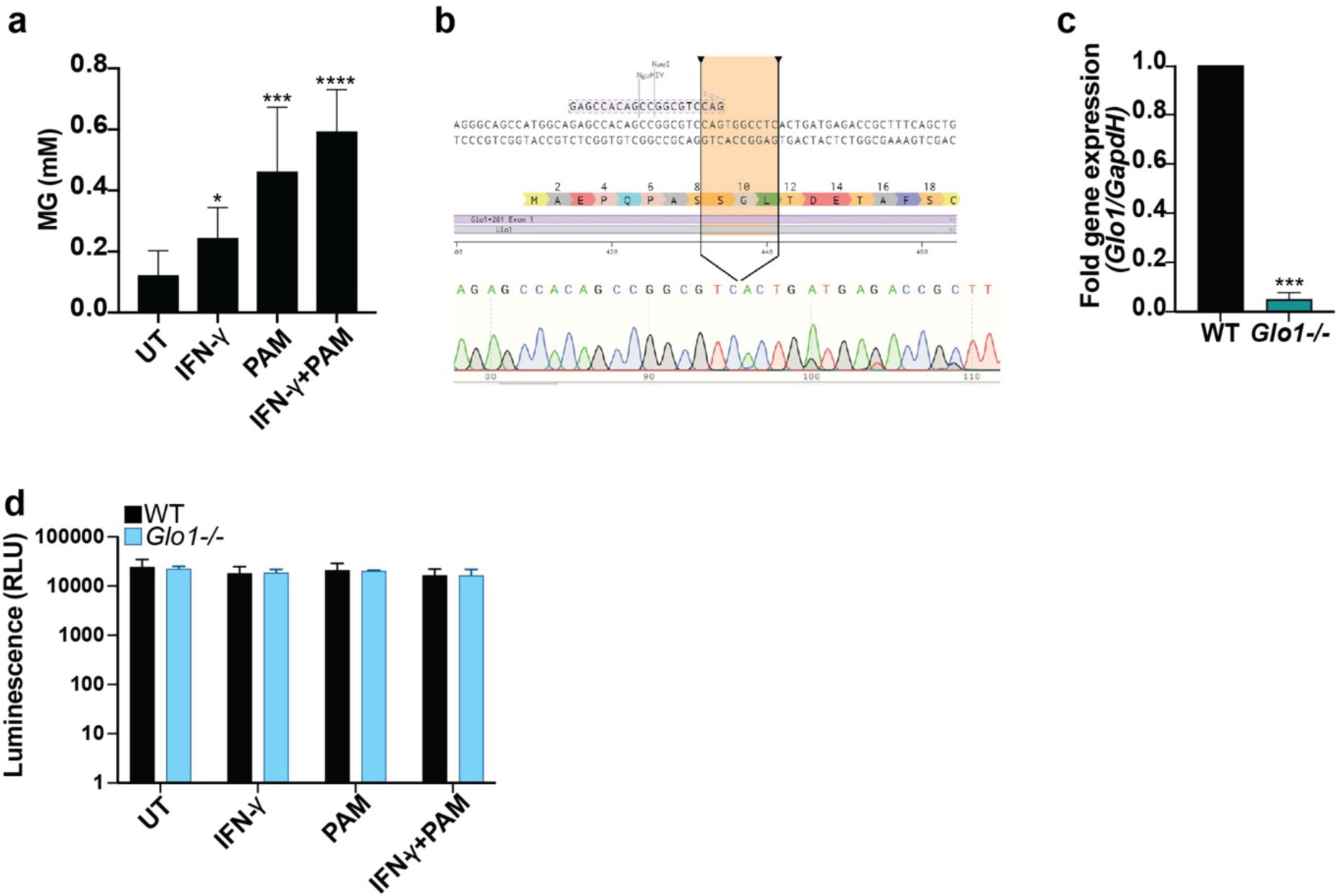
**(A)** Methylglyoxal measurements using a fluorogenic probe in uninfected wild-type B6 macrophages that were stimulated for 24-hours with IFN-γ or PAM3CSK4 (PAM) or both. **(B)** Sequencing of *Glo1−/−* mice that have a 10bp deletion resulting from the removal of nucleotides 432-441 from exon 1 of the *Glo1* gene. **(C)** qPCR data showing fold induction of *Glo1* in Wild-type and *Glo1−/−* B6 murine macrophages. **(D)** Viability measurements using CellGLOTitr of murine macrophages that were stimulated for 24-hours with IFN-γ or PAM3CSK4 (PAM) or both. Data represent at least three independent experiments. *P* values were calculated using Mann-Whitney test; \**P* < 0.05, \*\**P* < 0.01, \*\*\**P* < 0.001, **** indicates *P* <0.0001.

**Extended Data Figure 2.**
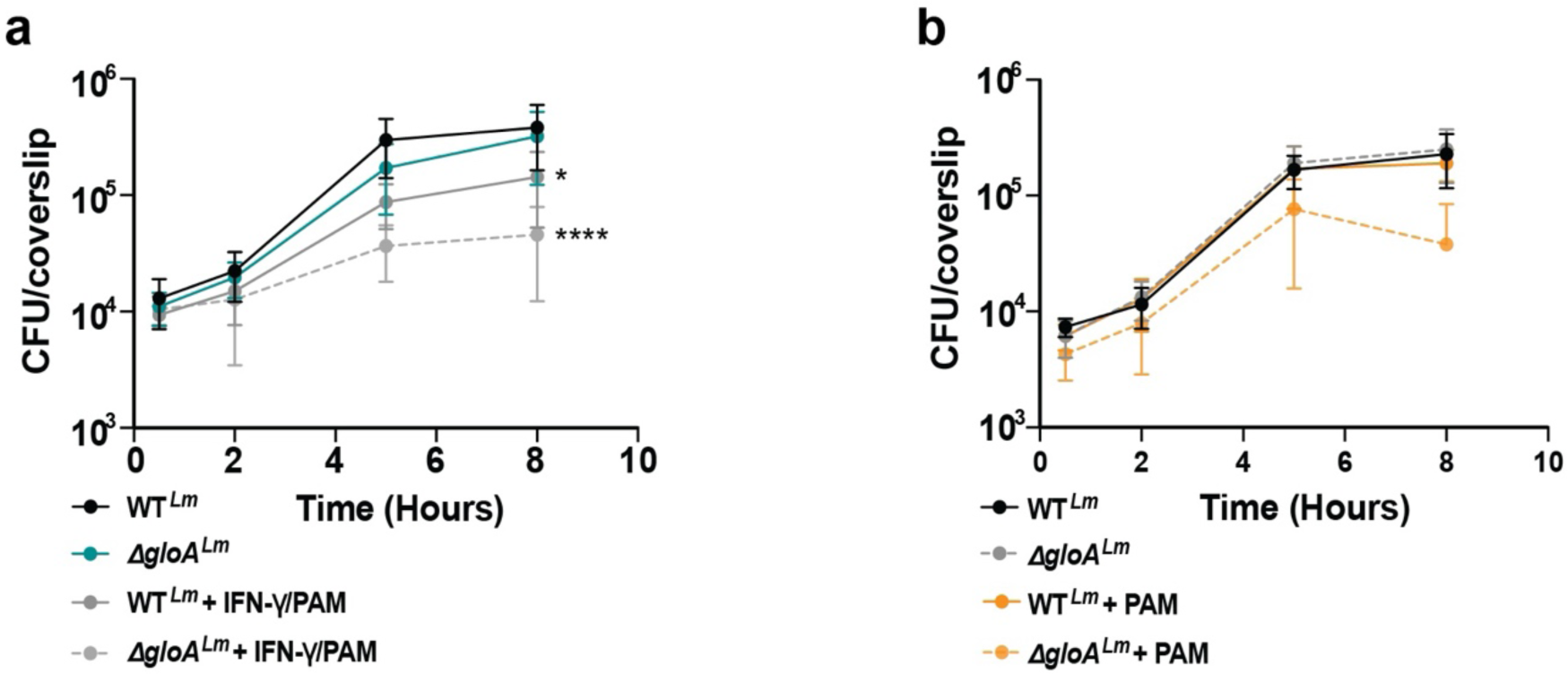
Wild-type B6 murine BMDMs were infected with wild-type *L. monocytogenes* or the *gloA* mutant without treatment or treated with IFN-γ **(A)** or PAM3CSK4 (PAM) **(B)** and intracellular CFU were enumerated at different time-points post-infection. Data are mean and SEM of three technical replicates of three independent experiments. Statistical significance is shown for *ΔgloA* infections compared to wild-type *L. monocytogenes* growing in wild-type macrophages. *P* values were calculated using Mann-Whitney test; \*\**P* < 0.05, \*\**P* < 0.01, \*\*\**P* < 0.001, **** indicates *P* <0.0001.

**Extended Data Figure 3.**
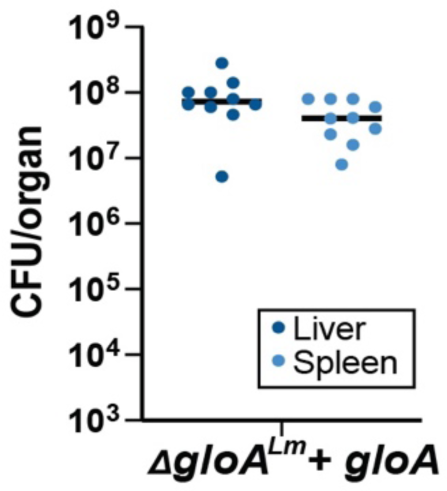
Wild-type B6 mice were infected with 10^5^ CFU of the *ΔgloA* complemented *L. monocytogenes* mutant. Livers and spleens were harvested 48-hours post-infection and CFU were enumerated. Data and median represent at least three pooled experiments.

**Extended Data Figure 4.**
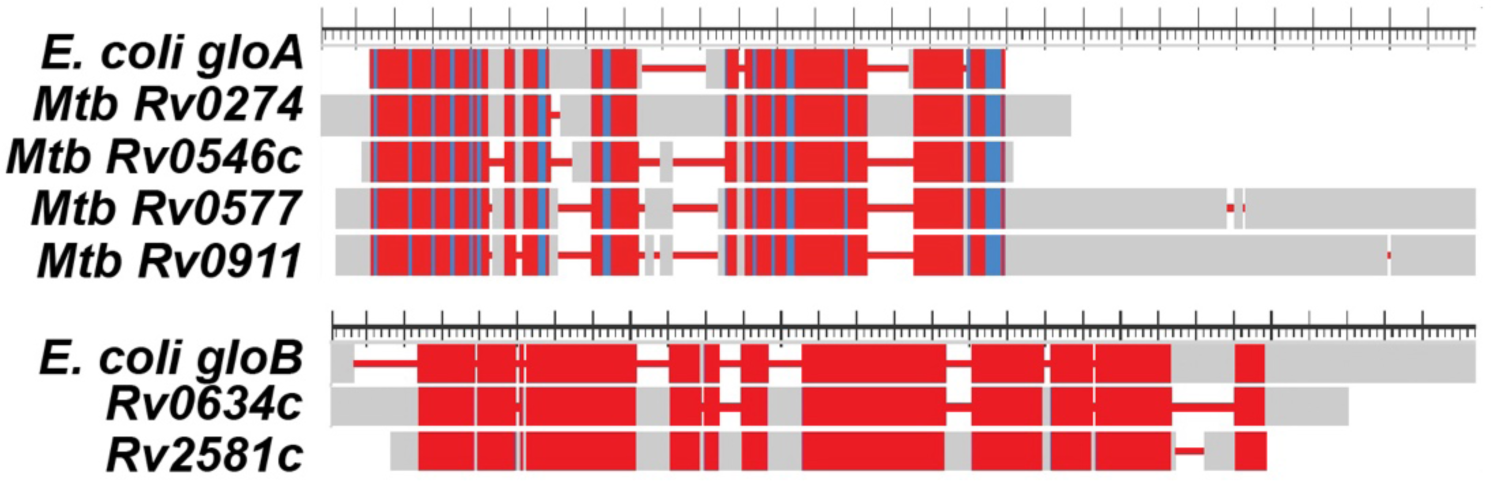
Protein sequence alignments of putative *Mtb* glyoxalases with *E. coli gloA or gloB.* Red indicates highly conserved residues, blue less conserved.

**Extended Data Figure 5.**
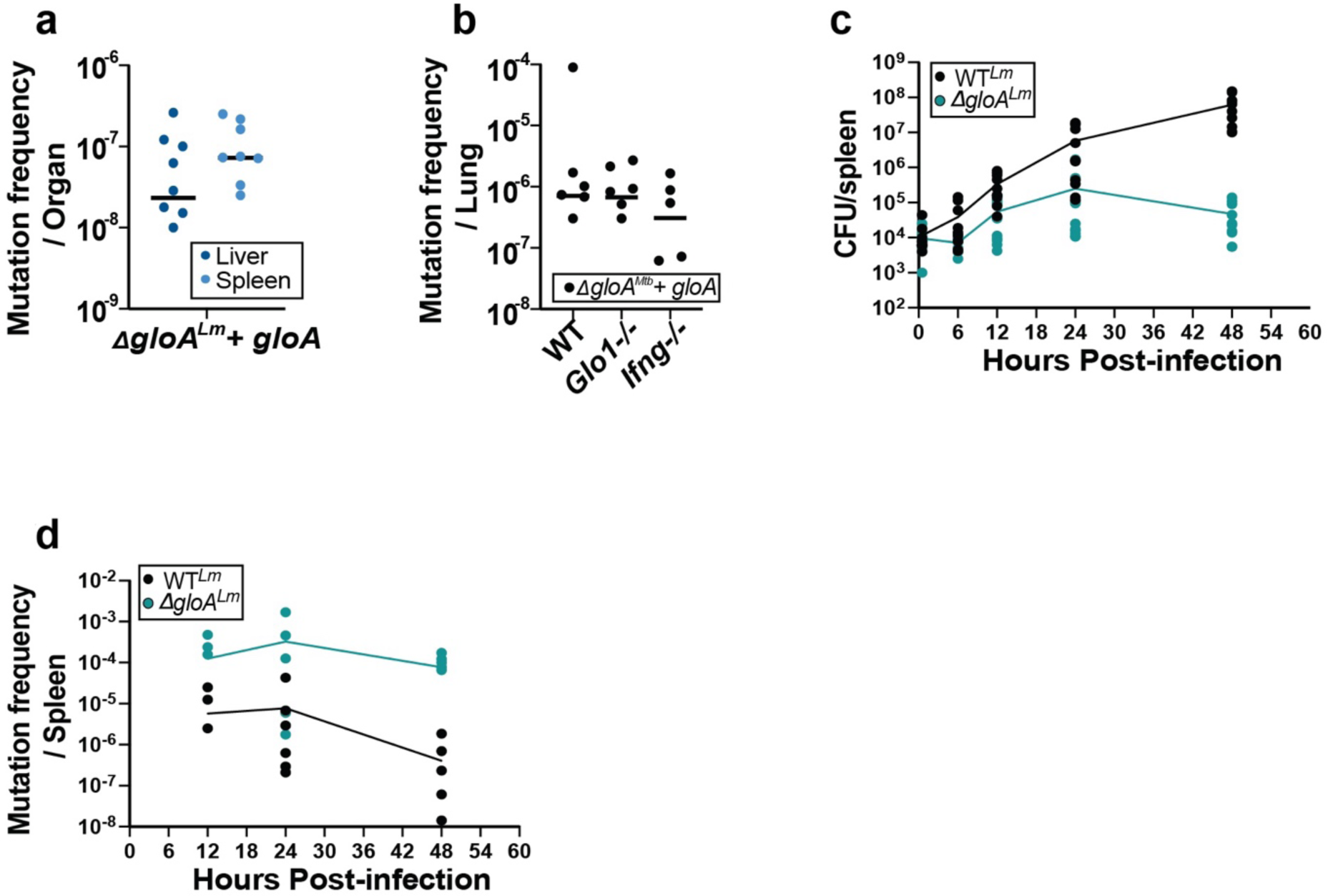
**(A)** Wild-type B6 mice were infected with 10^5^ CFU of the *ΔgloA* complemented *L. monocytogenes* mutant. Livers and spleens were harvested 48-hours post-infection and whole organs were plated in agar plates containing Rifampicin. **(B)** Wild-type B6 mice were aerosol-infected with 250 CFU of wild-type (H37Rv), Rv0911::tn or the complemented strain. Lungs were harvested at twenty-eight days post-infection and whole organs were plated on agar plates containing Rifampicin. **(C** and **D)** Wild-type B6 mice were infected with 10^5^ CFU of wild-type or *ΔgloA L. monocytogenes.* Spleens were harvested at various time-points post-infection and CFU **(C)** or mutation frequency **(D)** were determined for whole organs. For all experiments, mutation frequency was calculated as the total Rifampicin resistant colonies normalized to total CFU per organ. Data and median represent at least three independent experiments. *P* values were calculated using Mann-Whitney test; \**P* < 0.05, \*\**P* < 0.01, \*\*\**P* < 0.001, **** indicates *P* <0.0001.

**Extended Data Table 1.** Strains used in this study.

| Strain name | Description | Bacteria | Reference |
| --- | --- | --- | --- |
| 10403S | Wt | *L. monocytogenes* | (1) |
| DP-L7333 | $\Delta gloA$ | *L. monocytogenes* | (2) |
| DP-L7337 | $\Delta gloA$ pPL2.phyper.gloA | *L. monocytogenes* | (2) |
| DP-L7676 | $\Delta mgsA$ | *L. monocytogenes* | This study |
| DP-L7677 | $\Delta gloA \Delta mgsA$ | *L. monocytogenes* | This study |
| DP-E7678 | PLIM-$\Delta mgsA$ | *E. coli* SM10 | This study |
| XL 1 Blue | For vector construction | *E. coli* | Stratagene |
| SM10 | For conjugation | *E. coli* | (3) |
| H37Rv | Wt | *M. tuberculosis* |  |
| AAS-M1 | *Rv0911::tn* | *M. tuberculosis* | This study |
| AAS-M2 | *Rv0911::tn* PMV306-pNative-*Rv0911* | *M. tuberculosis* | This study |
| DP-E4073 | *Escherichia coli* K12 | *E. coli* | (4) |
| DP-E6631 | *Enterococcus durans* ATCC6056 | *E. durans* | (4) |
| DP-E6632 | *Enterococcus faecalis* ATCC 19433 | *E. faecalis* | (4) |
| DP-E6641 | *Bacillus subtilis* 168 | *B. subtilis* | (4) |
| U11 | *Francisella novicida* | *F. novicida* | (5) |
| ATCC LVS | Live-vaccine strain | *F. tularensis* | (6) |
| NR-46543 | JE2 USA300 Methicillin-resistant *Staphylococcus aureus* | MRSA | Bei resources |

1. Bécavin C, Bouchier C, Lechat P, Archambaud C, Creno S, Gouin E, et al. Comparison of Widely Used *Listeria monocytogenes* Strains EGD, 10403S, and EGD-e Highlights Genomic Differences Underlying Variations in Pathogenicity. *mBio*. 2014 May;5(2). 2. Anaya-Sanchez A, Feng Y, Berude JC, Portnoy DA. Detoxification of methylglyoxal by the glyoxalase system is required for glutathione availability and virulence activation in *Listeria monocytogenes*. *PLoS Pathog*. 2021 Aug 18;17(8):e1009819. 3. Simon R, Priefer U, Pühler A. A Broad Host Range Mobilization System for In Vivo Genetic Engineering: Transposon Mutagenesis in Gram Negative Bacteria. *Bio/Technology*. 1983 Nov;1(9):784–91. 4. Light SH, Su L, Rivera-Lugo R, Cornejo JA, Louie A, Iavarone AT, et al. A flavin-based extracellular electron transfer mechanism in diverse Gram-positive bacteria. *Nature*. 2018 Oct 12;562(7725):140–4. 5. Rohmer, L., Fong, C., Abmayr, S. et al. Comparison of *Francisella tularensis* genomes reveals evolutionary events associated with the emergence of human pathogenic strains. *Genome Biol* 8, R102 (2007). https://doi.org/10.1186/gb-2007-8-6-r102 6. Kurtz SL, Voskanian-Kordi A, Simonyan V, Elkins KL. Sequence comparison of *Francisella tularensis* LVS, LVS-G and LVS-R. *Pathog Dis*. 2018 Oct 1;76(7). doi: 10.1093/femspd/fty067. PMID: 30137434.

